# Revised annotation and characterization of novel *Aedes albopictus* miRNAs and their potential functions in dengue virus infection

**DOI:** 10.1101/2020.03.01.972398

**Authors:** Azali Azlan, Muhammad Amir Yunus, Ghows Azzam

## Abstract

The Asian tiger mosquito, *Aedes albopictus* (*Ae. albopictus*), is a highly invasive species that transmit several arboviruses including dengue (DENV), Zika (ZIKV), and chikungunya (CHIKV). Although several studies have identified microRNAs (miRNAs) in *Ae. albopictus*, it is crucial to extend and improve current annotations with the newly improved genome assembly, and the increase number of small RNA-sequencing data. We combined our high-depth sequence data and 26 public datasets to re-annotate *Ae. albopictus* miRNAs, and found a total of 110 novel mature miRNAs. We discovered that the expression of novel miRNAs was lower than known miRNAs. Furthermore, compared to known miRNAs, novel miRNAs are prone to be expressed in stage-specific manner. Upon DENV infection, a total of 59 novel miRNAs were differentially expressed, and target prediction analysis revealed that miRNA-target genes were involved in lipid metabolism and protein processing in endoplasmic reticulum. Taken together, miRNA annotation profile provided here is the most comprehensive to date, and we believed that this will facilitate future research in understanding virus-host interactions particularly on the role of miRNAs.

## Introduction

*Aedes albopictus* (*Ae. albopictus*) is an important vector of arboviruses including dengue (DENV), Zika (ZIKV) and chikungunya (CHIKV) (Chen et al., 2015). This mosquito is a robust and highly invasive species in both temperate and tropical regions of the world (Bonizzoni et al., 2013). Due to its high adaptability, *Ae. albopictus* has spread rapidly and is considered a public threat throughout the world; hence, studying the fundamental biology of this mosquito is important for controlling its aggressive spread.

Previous studies in *Ae. albopictus* have shown the importance of non-coding RNAs (ncRNAs) in development and virus infection (Batz et al., 2017; Liu et al., 2015; Su et al., 2019, 2017). Regulatory ncRNAs in metazoans include small RNAs, which consist of micro-RNAs (miRNAs), short-interfering RNAs (siRNAs), and PIWI-interacting RNAs (piRNAs). Particularly, miRNAs are 18-22 nucleotides (nts) in length, and function in the regulation of gene expression by targeting messenger RNAs (mRNAs), resulting in the post-transcriptional silencing of the target genes (Azlan et al., 2016; Coller and Parker, 2005; Djuranovic et al., 2012; Guo et al., 2010). Furthermore, miRNAs have been shown to play important roles in development, growth, and infection of animals (Bartel, 2009). For instance, through next-generation sequencing approach, multiple studies have reported that the expression profiles of *Ae. albopictus* miRNAs were altered at different developmental stages, during virus infection, and at diapause condition (Avila-Bonilla et al., 2017; Batz et al., 2017; Liu et al., 2015; Su et al., 2019, 2017). These studies suggest the involvement of miRNAs in the regulation of *Ae. albopictus* development and virus infection.

High quality genome coupled with high-depth RNA-sequencing data are important for accurate and comprehensive annotation of coding and non-coding genes. Alignment of small RNA reads to the reference genome is an essential step to annotate miRNAs in a species. Several studies used genome of closely related species as proxy reference to facilitate the miRNA expression analysis (Etebari et al., 2013; Su et al., 2019, 2017). Two previous studies by Su et al. 2017 and Su et al. 2019 used *Ae. aegypti* miRNAs in miRBase (http://www.mirbase.org/) to evaluate the expression of *Ae. albopictus* miRNAs in the midgut when exposed to (DENV) infection (Su et al., 2019, 2017). Both of the studies reported novel *Ae. albopictus* miRNAs without mapping small RNA reads against *Ae. albopictus* genome. Although this approach does not largely affect the differential expression analysis of highly abundant miRNAs in non-model organisms (Etebari and Asgari, 2014), the level of error in annotation is unknown. By using genome of closely related species to discover miRNAs of another organism, we limit ourselves to only miRNAs that are conserved by both species. miRNA target prediction software such as miRDeep2 (Friedländer et al., 2012) will excise potential precursors from the genome of closely related species based on the alignment of small RNA reads. Therefore, the precursor sequences predicted, which are derived from the reference genome of closely related species, may or may not be the true precursors of the newly identified miRNAs.

In 2015, *Ae. albopictus* genome (AaloF1) was sequenced using Illumina sequencing platform, and this draft assembly was derived from the Foshan strain in China (Chen et al., 2015). Using the aforementioned genome version, Batz et al. 2017 reported a total of 162 miRNAs in *Ae. albopictus* – 152 of them are known miRNAs, while the remaining 10 are novel (Batz et al., 2017). However, in 2018, a newer genome version of *Ae. albopictus* was published, derived from the C6/36 cell line (Miller et al., 2018). To date, this is the most contiguous and complete *Ae. albopictus* assembly published and available (assembly: canu_80X_arrow2.2, strain: C6/36, VectorBase). Unlike AaloF1, canu_80X_arrow2.2 used PacBio long-read sequencing technology (Miller et al., 2018), giving it an advantage in gene annotation and discovery. With this improved reference genome, we now have the chance to re-annotate miRNAs using the updated genome assembly together with additional small RNA sequencing libraries.

We first sequenced the small RNAs from DENV1-infected C6/36 cells to investigate miRNA expression upon DENV1 infection in *Ae. albopictus*. Before identifying the list of differentially expressed miRNAs upon infection, we first re-annotated miRNAs in *Ae. albopictus* using the updated genome assembly. To gain a comprehensive annotation catalog, we combined our C6/36 cell-derived sequencing libraries with 26 *Ae. albopictus* small RNA-seq publicly available datasets. These datasets encompassed multiple stages of *Ae. albopictus* development including embryo, larvae, pupae, adult males, sugar-fed and blood-fed females. We also included sequencing libraries from Batz et al. 2017 which derived from diapause and non-diapause pharate larvae (Batz et al., 2017). Finally, we predicted target sites of differentially expressed miRNAs to gain insights into their regulatory roles of *Ae. albopictus* in virus infection.

## Results and Discussion

### RNA-sequencing libraries and read mapping

In this study, we generated high-depth small RNA-seq libraries which were derived from DENV1-infected and uninfected *Ae. albopictus* C6/36 cell lines. All reads generated in this study were deposited in Short Read Archive (SRA) with the accession number SRP193815. A total of 270 millions cleaned reads of 18-32 bp in length were acquired from our pooled set of six C6/36 small RNA-seq libraries (**Table 1**). These 18-32 bp raw reads were aligned to *Ae. albopictus* genome (canu_80X_arrow2.2, VectorBase), and the resulting alignment files were subjected to downstream miRNA prediction analysis. In general, more than 95% of clean reads aligned to the reference genome (**Table 1**)

**Table 1:** Mapping statistics of small RNA reads generated in this study

| Sample | Raw reads | Clean reads | 18-32 bp reads | Alignment rate (%) |
| --- | --- | --- | --- | --- |
| DENV replicate 1 | 78933646 | 76813221 | 51893885 | 95.73 |
| DENV replicate 2 | 77299313 | 74393164 | 50852534 | 96.02 |
| DENV replicate 3 | 55658966 | 54155383 | 42163819 | 95.29 |
| Uninfected replicate 1 | 54757609 | 53082234 | 41476616 | 96.62 |
| Uninfected replicate 2 | 52985076 | 51516847 | 41877287 | 95.78 |
| Uninfected replicate 3 | 58901861 | 57008973 | 41997868 | 95.31 |

To visualize the heterogeneity of mapped reads in each library, total number of small RNAs and their non-identical reads were plotted according to their size (**Figure 1**). Length distribution of total small RNAs revealed a major peak in 22 nts size, but this peak collapsed when the total reads were converted into small RNA tags (Figure 1). The collapsibility of the peaks indicates the existence of miRNA in the libraries since miRNAs are usually made up of relatively few identical sequences (Armisen et al., 2009). Unlike 22 nt peak, peaks correspond to longer small RNAs (24-32 nt) did not heavily collapsed when converted into tags, suggesting the presence of complex pool of small RNA population (Harding et al., 2014).

**Figure 1:**
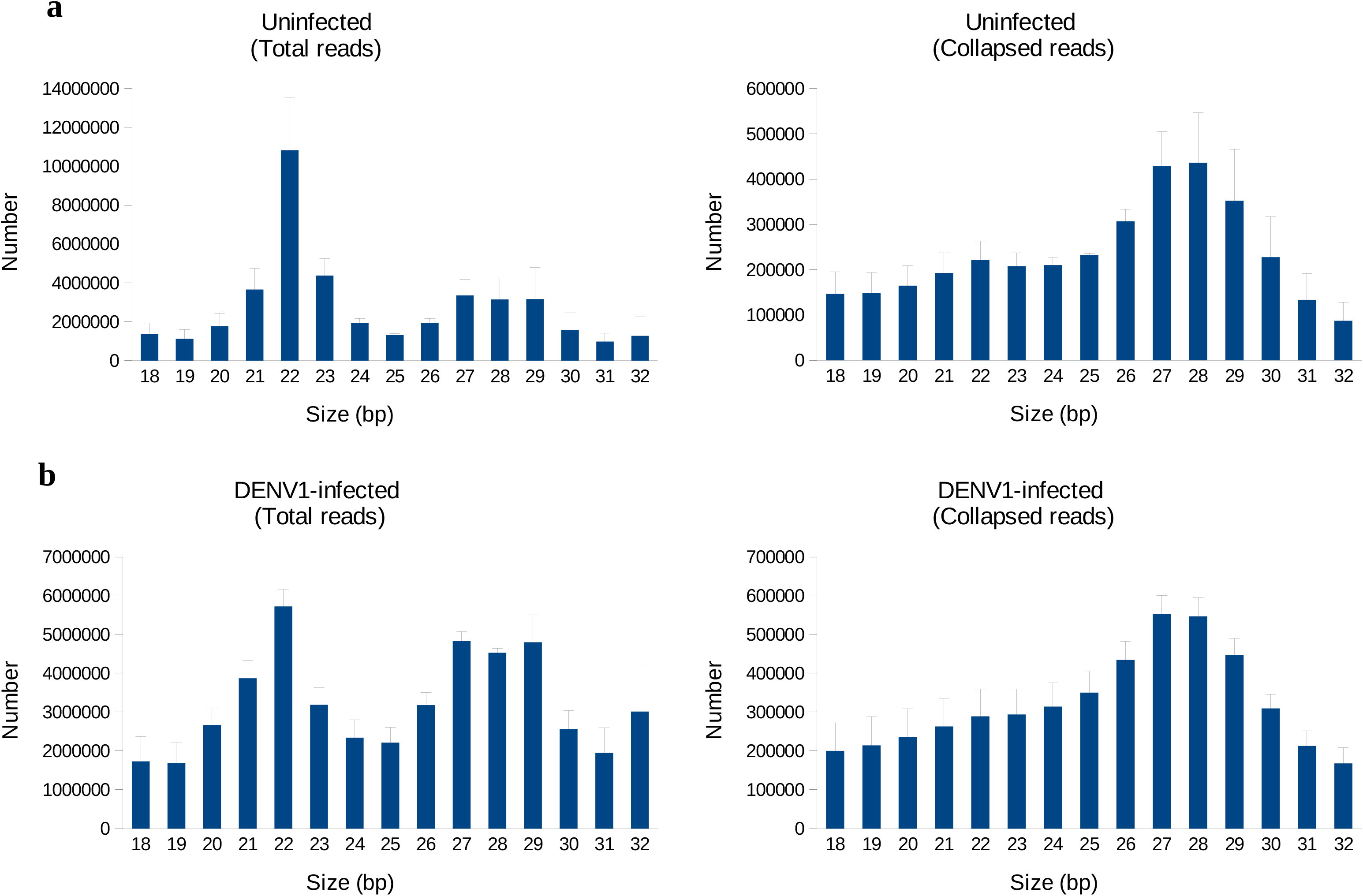
Length distribution of small RNAs in DENV1-infected and uninfected C6/36 cells. Size distribution of small RNA reads in uninfected samples (a) and DENV1-infected samples (b).

### Identification of novel miRNAs in *Ae. albopictus*

To identify putatively novel miRNAs for *Ae. albopictus*, we combined reads from 6 libraries generated in this project with 26 public datasets, resulting in a total of approximately 350 million reads. The reads were then mapped to the C6/36 genome (canu_80X_arrow2.2, VectorBase), and the resulting alignment file was fed into miRDeep2 for novel miRNA discovery. In our miRDeep2 analysis, we used miRNAs predicted from Batz et al. 2017 as known *Ae. albopictus* miRNAs. We also defined other known miRNAs as miRNAs that have sequence conservation with miRNAs from other species especially from the arthropod family curated from miRBase 2.1 (http://www.mirbase.org/). The majority of conserved miRNAs have been identified by previous studies (Avila-Bonilla et al., 2017; Batz et al., 2017; Gu et al., 2013; Su et al., 2019, 2017). We successfully confirmed the presence of miRNAs previously predicted by Batz et al. 2017 in canu_80X_arrow2.2 genome version (**Supplementary Info 1**) (Batz et al., 2017). A total of 110 novel mature miRNAs which derived from 125 precursors were identified. Complete list of both mature and precursor novel miRNAs can be found in Supplementary Info 1. The majority of mature *Ae. albopictus* miRNAs were found to be 22 bp in length, while in the case of precursor miRNAs, most of them ranged between 58-65 bp (**Figure 2**). *Ae. albopictus* miRNAs varied in terms of genomic loci – 49.9% and 48.2% of them located in the intergenic and intronic regions, while the remaining resided within coding sequence (CDS).

**Figure 2:**
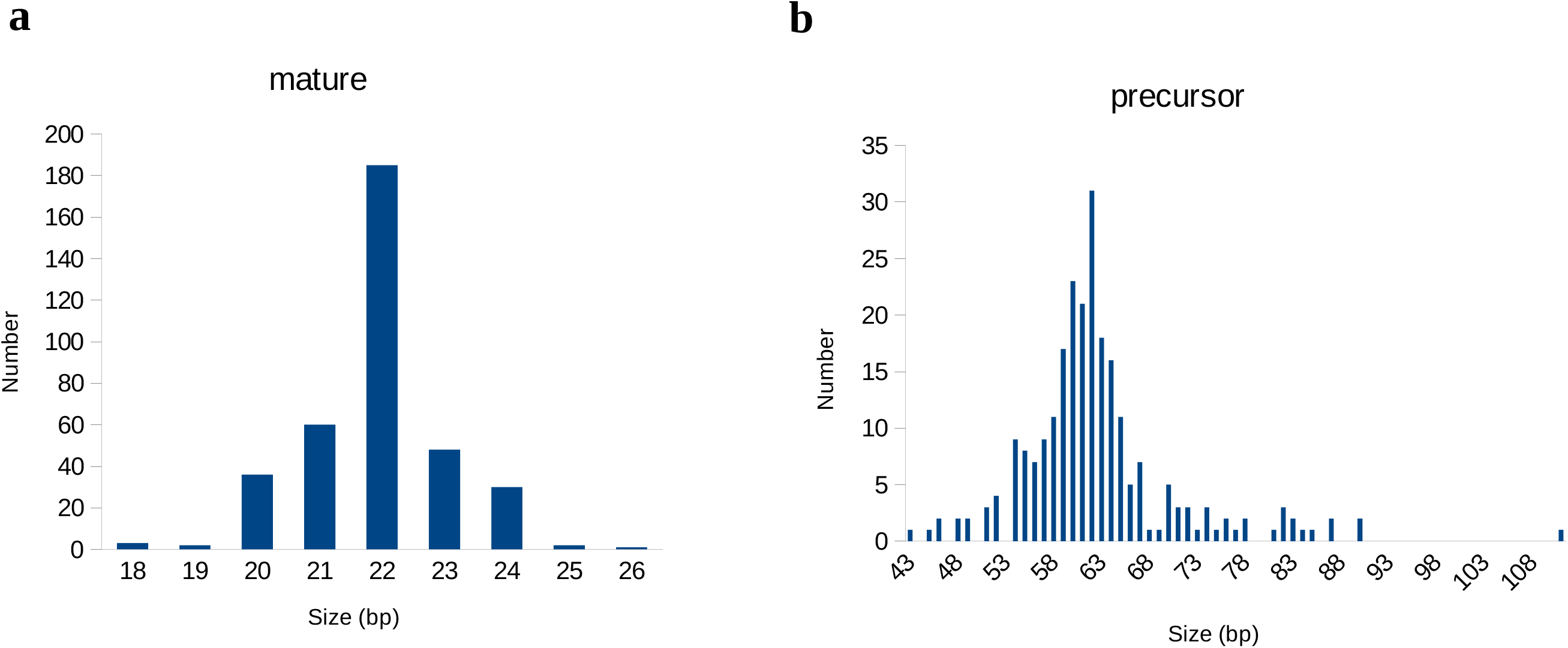
Size distribution of mature and precursor miRNAs. Size distribution of mature miRNA (a) and precursor miRNA (b)

We found that 35 of novel miRNAs shared the same seed sequence with arthropod miRNAs curated from miRBase 2.1 (http://www.mirbase.org/) (**Supplementary Info 1**). These novel miRNAs shared the same seed region, which, according to miRDeep2, is the first 18 nt of a read sequence (Friedländer et al., 2012). Despite sharing the same seed sequence, these miRNAs were still considered novel because the predicted precursor sequences were not similar to known precursors. miRNAs interact with their target mRNA via base pairing of its “seed” sequence, which usually in the position of 2-8 from the 5’ end of the mature miRNAs (Lai, 2002). Hence, these 35 novel miRNAs may target the same genes as the known miRNAs which they shared the same seed region with.

The number of novel miRNAs detected in this study was higher compared to other *Ae. albopictus* studies (Batz et al., 2017; Gu et al., 2013; Liu et al., 2015). For example, Gu et al., 2013 reported the discovery of 15 novel miRNAs, whereas Liu et al., 2015 and Batz et al. 2017 identified 5 and 10 novel miRNAs in *Ae. albopictus* respectively. This finding is presumably due to high sequencing depth and high percentage of mapped reads in every replicate of C6/36 small RNA libraries generated in this study. It was reported that libraries with lower number of reads make the discovery of miRNAs, especially the lowly expressed, more challenging and somewhat limited (Metpally et al., 2013). Another plausible explanation for this high number of novel miRNAs is the large genome size of *Ae. albopictus*; in fact, it is the largest among mosquitoes genomes ever sequenced. Genome size of Foshan (AaloF1) and C6/36 (canu_80X_arrow2.2) assemblies are 2,247 Mb and 1,967 Mb respectively (Chen et al., 2015; Miller et al., 2018), while *Ae. aegypti* is 1,279 Mb (Matthews et al., 2018), *Culex quinquefasciatus (C. quinquefasciatus)*, and *Anopheles darlingi (An. darlingi)* are 540 Mb and 174 Mb respectively (Arensburger et al., 2010; Holt et al., 2002).

We also discovered that multiple locations in *Ae. albopictus* genome can give rise to the same mature and precursor miRNAs. A total of 168 distinct locations were found to be the genomic loci of 120 novel miRNAs, suggesting gene duplication event. This was also true for *Ae. albopictus* protein-coding genes. Duplication of several members of gene families involved in insecticide-resistance, diapause, immunity, olfaction, and sex determination was observed in *Ae. albopictus* genome, and this contribute to its large genome size (Chen et al., 2015).

From the standpoint of miRNA biogenesis, generation of functional novel mature miRNAs requires the transcription of certain genomic loci that are capable of producing RNA whose secondary structures are recognizable by Drosha and Pasha (DGCR8 in vertebrates) in the nucleus, and Dicer and Loquacious in the cytoplasm (Azlan et al., 2016; Denli et al., 2004; Gregory et al., 2004; Lee et al., 2003). Therefore, the birth of novel miRNA requires relatively simple prerequisite, which is loci in the genome that evolve to produce RNAs that fold in such a way that miRNA protein machineries can identify and process them, resulting in fully functional mature miRNAs (Berezikov, 2011; Chen and Rajewsky, 2007) Due to inherent property of RNA to form imperfect folding (biogenesis of miRNAs requires imperfect folding of RNA hairpins), the emergence of novel miRNAs, from the evolutionary viewpoint, is prone to be more feasible than the birth of new coding genes (Chen and Rajewsky, 2007).

### Expression of novel miRNAs in different stages of *Ae. albopictus* development

We then assessed the dynamics of novel miRNA expression throughout *Ae. albopictus* development to have a more understanding of its distribution. We used public dataset from Gu et al. 2013 which consists of six different stages namely, 0-24 hour embryo, larvae, pupae, adult males, blood-fed females and sugar-fed females (Gu et al., 2013). In general, the expression level of novel miRNAs was lower than known miRNAs across all six stages (**Figure 3a**). Next, we determined the specificity of novel miRNA expression across different developmental stages. We calculated the specificity score of each miRNA using Jensen-Shannon (JS) score, which ranges from zero to one (Cabili et al., 2011). This specificity metric quantifies the similarity in expression value across developmental stages. miRNAs having JS score of one indicates the extreme case in which it is only expressed in one specific stage, whereas those having a score close to zero are ubiquitously expressed in all stages with relatively similar value of expression (Cabili et al., 2011).

**Figure 3:**
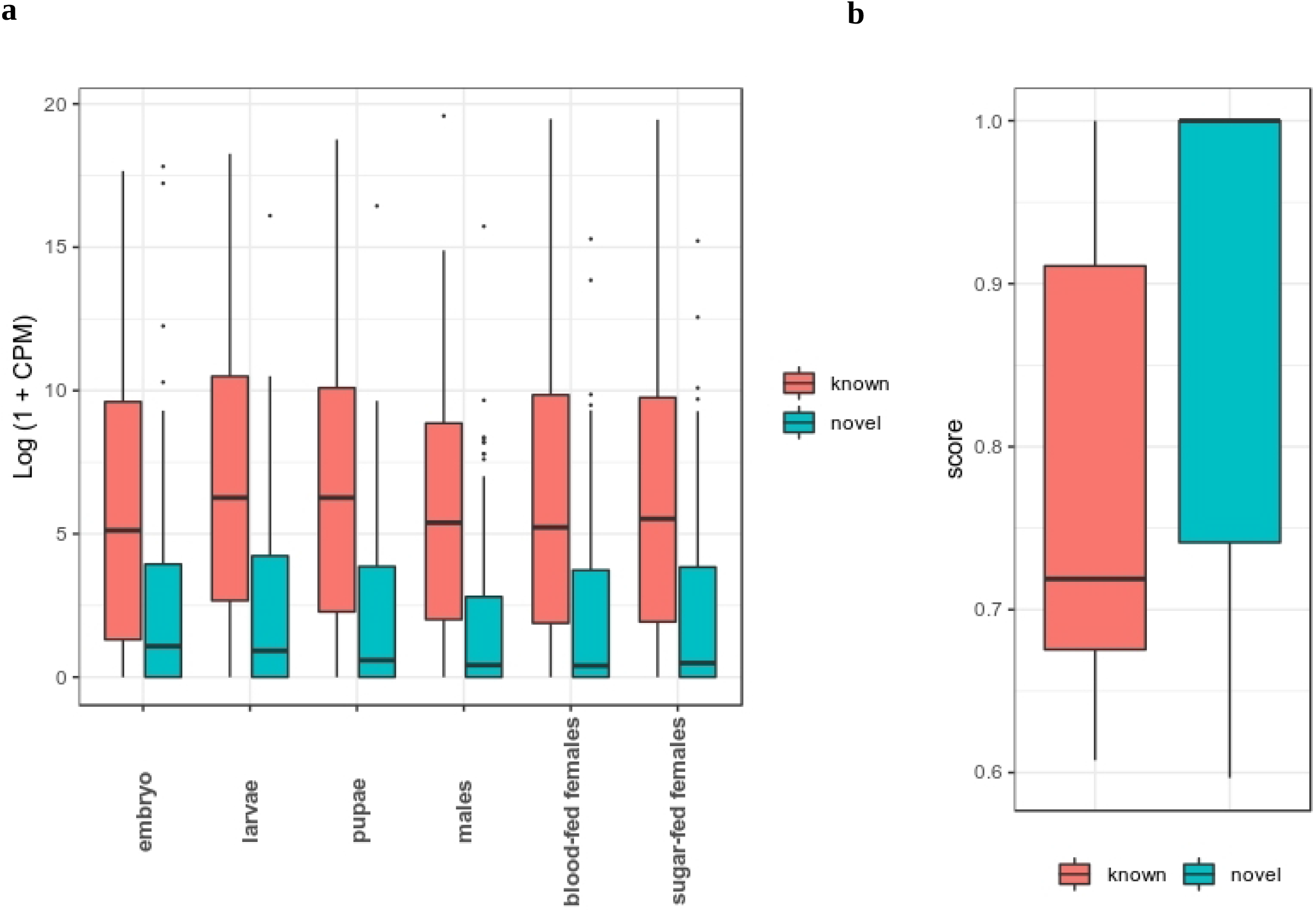
Expression distribution and JS specificity score of known and novel miRNAs. (a) Expression distribution of lncRNAs and protein-coding genes in *Cx. quinquefasciatus* (b) Distribution of specificity score of lncRNAs and protein-coding genes in *Cx. quinquefasciatus*

**Figure 4:**
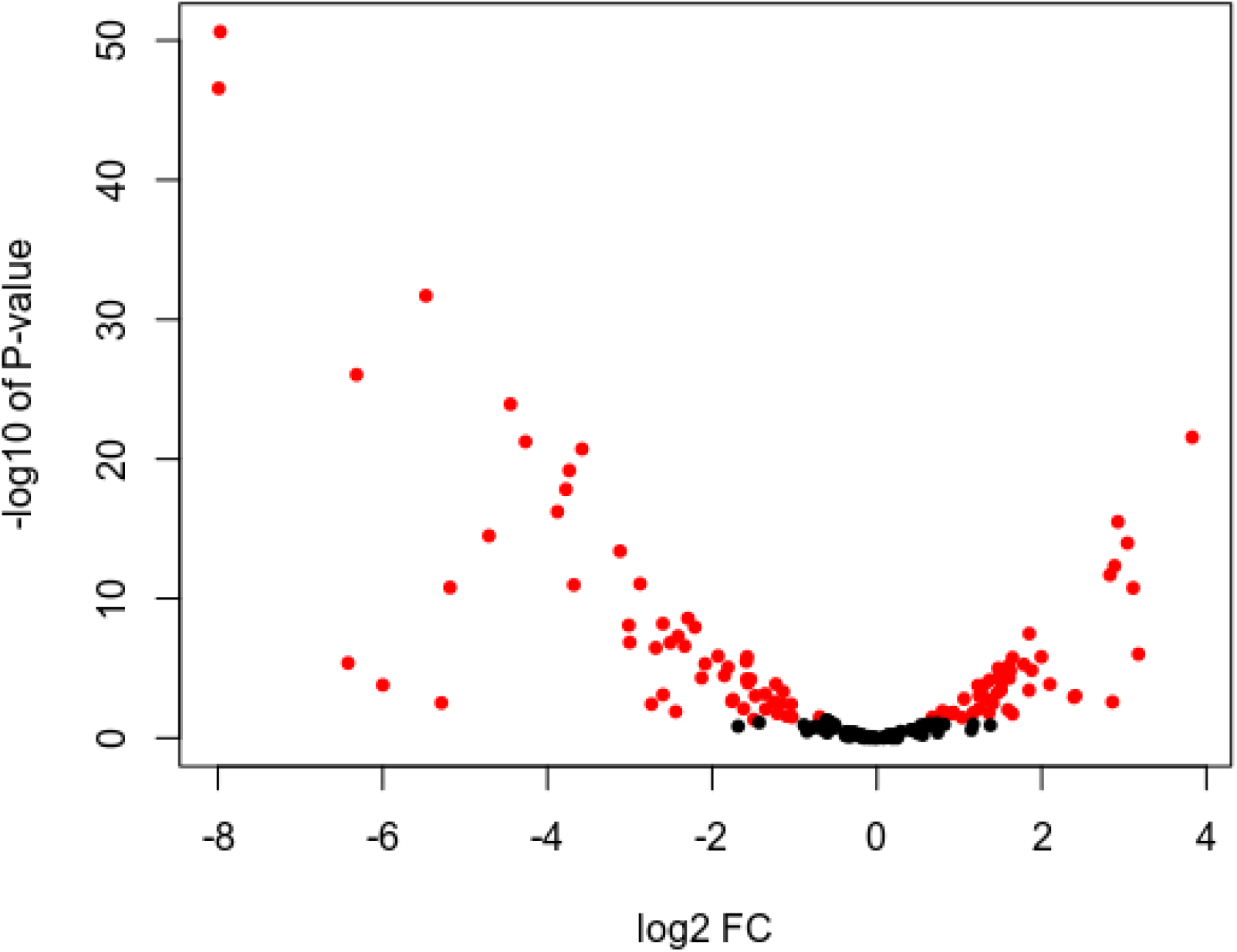
Volcano plot of differentially expressed miRNAs upon DENV1 infection. Red dots represent significant P-value (<0.05), while black dots represent insignificant P-value (>0.05)

Here, we discovered that novel miRNAs were more specific in expression across different developmental stages compared to known miRNAs (**Figure 3b**). For instance, 23% of known miRNAs had JS score of 1, while for novel miRNAs, 46% of them had JS score of 1. Known miRNAs, on the other hand, have a more ubiquitous expression profile. This observation indicates that, unlike known miRNAs, novel miRNAs possess a narrower window in expression than known miRNAs. We speculate that novel miRNAs in *Ae. albopictus* possibly function as stage-specific tuners and regulators of genes involved in the control of developmental transition. Despite their lower expression, the specificity of novel miRNAs expression hinted to the possibility that they may execute distinct functions at specific developmental time-point.

### *Ae. albopictus* novel miRNAs were differentially expressed upon DENV infection

Previous studies have reported that *Ae. albopictus* miRNAs experienced changes in expression level upon DENV infection (Avila-Bonilla et al., 2017; Liu et al., 2015; Su et al., 2019, 2017). We asked if our newly annotated miRNAs were differentially expressed upon virus infection. We chose DENV serotype 1 (DENV1) in this study. DENV serotype 2 (DENV2) have been used in many genome-wide expression studies either in mammalian or mosquito hosts (Angleró-Rodríguez et al., 2017; Bonizzoni et al., 2012; Miesen et al., 2016; Sim et al., 2013; Sim and Dimopoulos, 2010; Tsujimoto et al., 2017; Zanini et al., 2018). DENV2 has been the center of attention in studies involving DENV, despite the fact that other serotypes are similarly as important as serotype 2. Different serotype of DENV display different clinical manifestation and level of severity. A case study in Singapore reported that patients with DENV1 infection may exhibit more severe illnesses compared to those infected with DENV2 (Yung et al., 2015). Besides, different symptoms were observed between DENV1 and DENV2-infected patients (Yung et al., 2015), suggesting that different serotype evokes different molecular responses in humans. Therefore, we infer that different serotype of DENV will elicit different transcriptional responses in *Ae. aegypti* mosquitoes.

In this study, we discovered that our newly annotated *Ae. albopictus* miRNAs in C6/36 cells were differentially expressed after 3 days post-infection with DENV1. A total of 102 miRNAs were found to be differentially expressed (P-value <0.05), whereby 59 of them were novel miRNAs. To access our small RNA sequencing data, five novel miRNAs that were differentially expressed were further validated using qRT-PCR (**Figure 5**).

**Figure 5:**
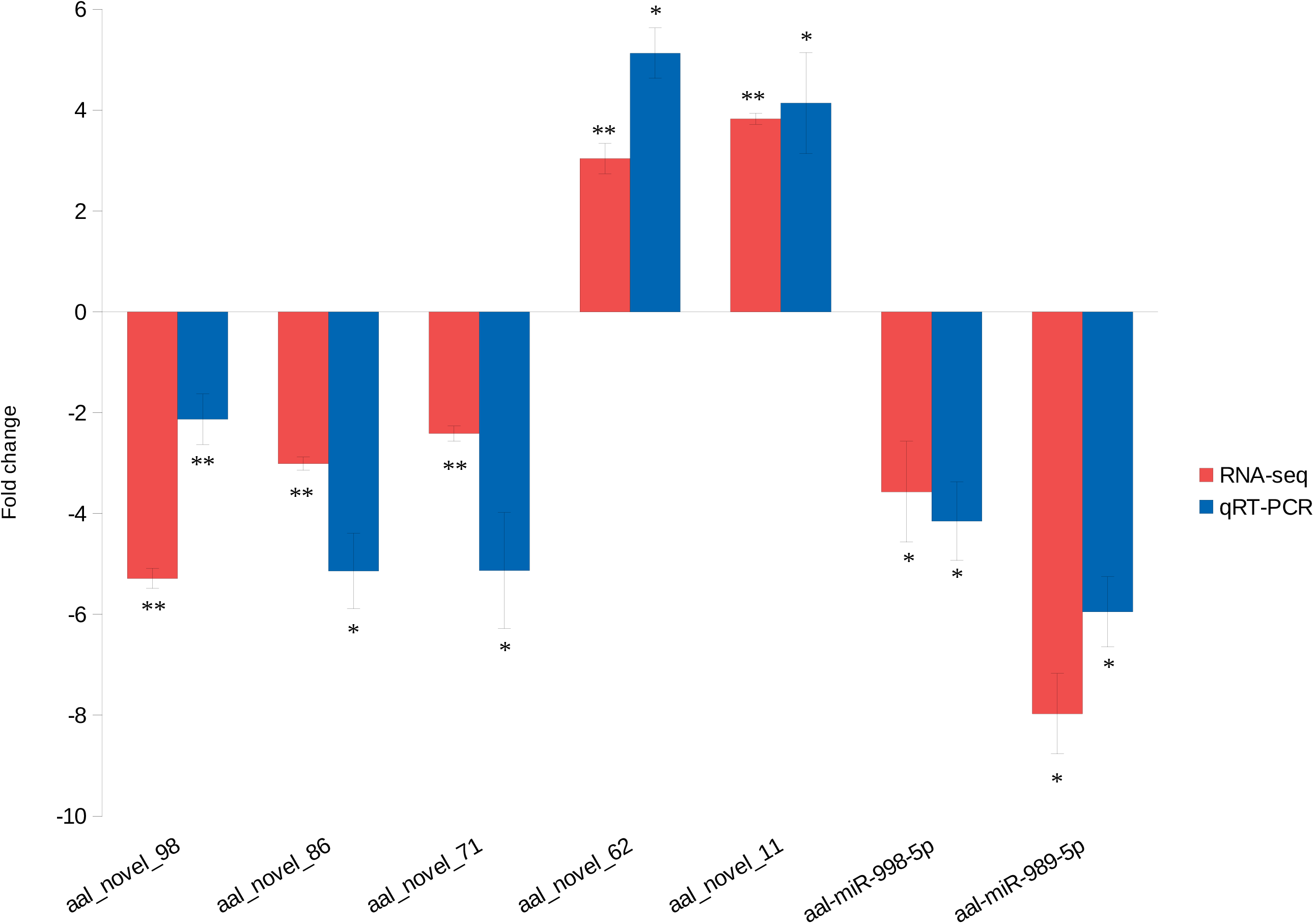
qRT-PCR of selected novel miRNAs. Two-tailed *t-*test was used for all comparisons; *p < 0.05, **p < 0.01. Expression levels were normalised to uninfected samples. Error bars indicate standard error of the mean (SEM) of three independent experiments.

Avila-Bonilla et al. 2017 performed small RNA-sequencing of C6/36 cells that were persistently infected with DENV2 to analyze changes in miRNA expression profile (Avila-Bonilla et al., 2017). Several differentially expressed miRNAs in C6/36 cells persistently infected with DENV2, such as miR-190, miR-970, miR-263a, miR-92b, and miR-9a-5p (Avila-Bonilla et al., 2017), were also found to be differentially expressed in our DENV1-infected C6/36 cells. Therefore, we hypothesized that these miRNAs may involve in host-DENV interaction in *Ae. albopictus*, regardless of the virus serotype. Meanwhile, other miRNAs may participate in the serotype-specific antiviral immunity in *Ae. albopictus*.

Next, we performed miRNA target prediction to gain further insights into the potential downstream effects of regulation by differentially expressed novel miRNAs in C6/36 cells following DENV 1 infection. Undeniably, identification of miRNA target genes remains a complicated task. *In silico* analysis and computational methods are important in predicting potential miRNA target genes, despite the outcomes of such approaches can vary from one algorithm to another (Etebari and Asgari, 2016). To alleviate the certainty in the miRNA target prediction, we merged the outcomes from two miRNA target prediction tools, namely miRanda (Enright et al., 2003) and RNAhybrid (Krüger and Rehmsmeier, 2006). miRNA binding sites derived from both softwares were deemed to be highly confident.

Both miRANDA and RNAhybrid predicted a total of 10,867 genes as targets for differentially expressed miRNAs. Gene Ontology (GO) analyses using g:Profiler (Reimand et al., 2007) showed that these target genes were involved in many types of biological process, and molecular functions inside cells such as DNA and protein-binding, transport and localization (**Figure 6**). Meanwhile, KEGG pathway analysis revealed two important pathways: glycerophospholipid metabolism and protein processing in endoplasmic reticulum (ER). Previous studies have reported that among the cellular factors required for the life cycle of enveloped virus, including DENV, are lipids. DENV utilized a specific lipid, bis(monoacylglycero)phosphate, which functions as a co-factor for viral fusion in late endosomes (Zaitseva et al., 2010). DENV entry into host cells is established via endocytosis and use lipids as factors that triggers the fusion of viral particles into endosome, hence, delivering the viral genome into cytosol (Mukhopadhyay et al., 2005; Rodenhuis-Zybert et al., 2010). One of the critical steps in DENV life cycle is the translation of viral genome into a single polypeptide chain by host ribosomes, and this process occurs in rough endoplasmic reticulum (Mukhopadhyay et al., 2005; Rodenhuis-Zybert et al., 2010). Therefore, post-transcriptional regulation of proteins involved in lipid metabolism and protein processing in ER may be crucial in either facilitating or inhibiting virus replication.

**Figure 6:**
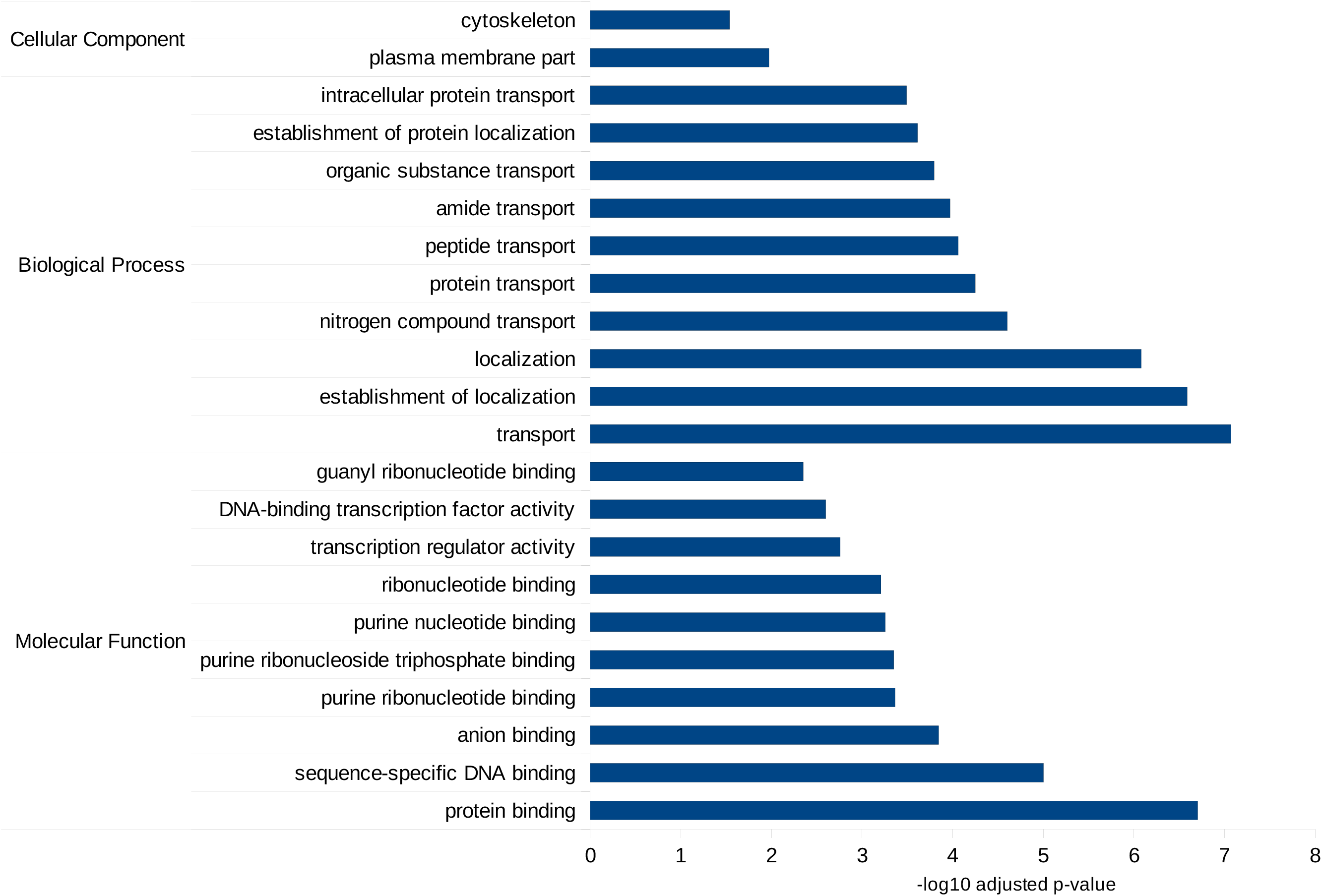
GO bar graphs of target genes of the differentially expressed miRNAs. Only top 10 of GO terms were plotted in the bar graphs.

## Experimental Procedures

### Cell culture and virus

*Ae. albopictus* C6/36 cell (ATCC: CRL-1660) were cultured in Leibovitz’s L-15 medium (Gibco, 41300039), supplemented with 10% Fetal Bovine Serum (FBS, Gibco, 10270) and 10% Tryptose Phosphate Broth (TPB) solution (Sigma, T9157). The C6/36 cells were incubated at 25°C without CO_2_.

BHK-21 cells (ATCC: CCL-10) were cultured at 37°C in Dulbecco’s modified Eagles Medium (DMEM, Gibco, 11995065) supplemented with 10% FBS (Gibco, 10270), and 5% CO_2_. Dengue virus serotype 1 (Hawaiian strain) was propagated in C6/36 cells and titered using BHK-21 cells. Determination of DENV1 titer was done using 50% tissue culture infectious dose – cytopathic effect (TCID50-CPE) as previously described (Atieh et al., 2016; Li et al., 2011). DENV1 (Hawaiian strain) used in this study was a gift from David Perera, University Malaysia Sarawak.

### Virus infection, RNA extraction and sequencing

C6/36 cells were infected with DENV1 at multiplicity of infection (MOI) of 0.25. After 3 day post infection, RNA extraction was carried out using miRNeasy Mini Kit 50 (Qiagen, 217004) according to the manufacturer’s protocol. Total RNA was then subjected to next-generation sequencing. The RNA-sequencing libraries were prepared using standard Illumina protocols and sequenced using HiSeq-SE50 platform generating paired-end reads of 50 bp in size.

### Verification of DENV1 infection in C6/36 cells

To verify the presence of DENV1 infection in C6/36 cells, total RNA of both uninfected and DENV1-infected samples were subjected cDNA synthesis using Tetro cDNA synthesis kit (Bioline, BIO-65042). Reverse oligo of DENV1 specific primer was used in cDNA synthesis. Polymerase chain reaction (PCR) assay was used performed using DENV1 specific primers (Johnson et al., 2005).

### Preparation of public datasets

Publicly available small RNA-seq datasets were downloaded from NCBI Sequence Reads Archive (SRA). List of public datasets can be found in **Supplemental Table 1**. Prior to downstream analyses, small RNA-seq adapters were clipped using Trimmomatic version 0.38 (Bolger et al., 2014), and low quality reads were removed. Reads of 18-32 bp size were retained for downstream analysis.

### miRNA identification

To identify putatively novel miRNAs, clean reads from all small RNA-seq used in this study were pooled and aligned to the *Ae. albopictus* genome, (canu_80X_arrow2.2, VectorBase), using miRDeep2 (Friedländer et al., 2012). We used miRNAs predicted by Batz et al. 2017 as known miRNAs. For known miRNA precursors, we used all precursor miRNAs within arthropod family that were retrieved from miRBase2.1. Predicted precursors that had significant Randfold p-value (p-value < 0.05) were retained (Bonnet et al., 2004). Minimum miRDeep score was set to >4. Then, we retained predicted miRNAs that share the same seed sequence as arthropod miRNAs. For miRNAs that did not share same the seed sequence, only predicted precursors that had at least 1000 reads of mature miRNAs or those having at least 10 reads of predicted star sequence were retained.

### Differential expression of miRNA

Quantifier module in miRDeep2 package was used to quantify miRNAs in uninfected and DENV1-infected libraries (Friedländer et al., 2012). In brief, total small RNA reads of both uninfected and DENV1-infected libraries were aligned against precursor miRNAs, and the resulting number of mapped reads indicate the miRNA abundance within each sample. Raw counts generated from the quantifier module were subjected to further analysis in R/Bioconductor environment using edgeR package (Robinson et al., 2010)

### miRNA target prediction and functional analysis

Target prediction was performed using miRANDA (Enright et al., 2003) and RNAhybrid (Krüger and Rehmsmeier, 2006) using 3’UTR extracted from *Ae. albopictus* GTF annotation file (Vectorbase). miRANDA predictions were performed using the following parameters: score cut-off 140, energy cutoff −20, gap open penalty −9, gap extension penalty −4, seed regions at positions 2-8. RNAhybrid predictions were performed using the following parameters: energy cut-off −20, p-value < 0.1, binding required in miRNA positions 2-7. Functional analysis was conducted using g:Profiler (Reimand et al., 2007). We used g:SCS threshold for multiple testing correction.

### Quantitative real-time PCR (qRT-PCR)

cDNA synthesis was done using miScript II RT Kit (Qiagen: 218160) followed by qRT-PCR with miScript SYBR Green PCR Kit (Qiagen: 218073) according to manufacturer’s protocol. *Ae. albopictus* housekeeping genes *RPS17* was used as an internal control for qRT-PCR (Dzaki and Azzam, 2018; Su et al., 2017), and the 2^-ΔΔCT^ method was used to determine relative expression levels of miRNAs. The experiment was performed using Applied Biosystems Step One PlusTM Real-Time PCR System.

**Supplemental Info 1: List of known and novel miRNAs in *Ae. albopictus* genome-wide**

**Supplemental Table 1: List of public datasets used in this study**

**Supplemental Table 2: List of primers used in this study**

**Supplemental Table 3: List of differentially expressed miRNAs upon DENV1 infection in C6/36 cells**

**Supplemental Table 4: List of GO terms of target genes of the differentially expressed miRNAs**

