## Supplemental for "Revised annotation and characterization of novel *Aedes albopictus* miRNAs and their potential functions in dengue virus infection"

**Supplemental Table 1: List of public datasets used in this study**

| Accession number | Condition | Reference/Publication |
|---|---|---|
| SRR5168339 | Mature oocytes diapause replicate 1 | Batz et al., 2017 |
| SRR5168338 | Mature oocytes diapause replicate 2 | |
| SRR5168337 | Mature oocytes diapause replicate 3 | |
| SRR5168336 | Mature oocytes non-diapause replicate 1 | |
| SRR5168335 | Mature oocytes non-diapause replicate 2 | |
| SRR5168334 | Mature oocytes non-diapause replicate 3 | |
| SRR5168333 | Pharate larvae diapause replicate 1 | |
| SRR5168332 | Pharate larvae diapause replicate 2 | |
| SRR5168331 | Pharate larvae diapause replicate 3 | |
| SRR5168330 | Pharate larvae diapause replicate 4 | |
| SRR5168329 | Pharate larvae non-diapause replicate 1 | |
| SRR5168328 | Pharate larvae non-diapause replicate 2 | |
| SRR5168327 | Pharate larvae non-diapause replicate 3 | |
| SRR5168326 | Pharate larvae non-diapause replicate 4 | |
| SRR609263 | Sugar-fed females | Gu et al., 2013 |
| SRR609265 | Adult males | |
| SRR609264 | Pupae | |
| SRR609260 | Larvae | |
| SRR609261 | Blood-fed females | |
| SRR609439 | 0-24 hours embryo | |
| SRR5251234 | persistently infected DENV2 replicate 2 | Avila-Bonilla et al., 2017 |
| SRR5251235 | persistently infected DENV2 replicate 1 | |
| SRR5251236 | acutely infected DENV2 replicate 2 | |
| SRR5251237 | acutely infected DENV2 replicate 1 | |
| SRR5251238 | not infected replicate 2 | |
| SRR5251239 | not infected replicate 1 | |

**Supplemental Table 2: List of differentially expressed miRNAs upon DENV1 infection in C6/36 cells**

| miRNA | logFC | logCPM | PValue | FDR |
|---|---|---|---|---|
| aal-miR-989-5p | -7.97160685824 | 15.4006287012213 | 2.38304822292902E-51 | 4.43246969464798E-49 |
| aal-miR-278-5p | -7.99015158479353 | 8.71877971937138 | 2.71982766222985E-47 | 2.52943972587376E-45 |
| aal-miR-7-3p | -5.47459399815262 | 4.88707154454216 | 1.97642440358462E-32 | 1.22538313022247E-30 |
| aal-miR-263a-3p | -6.3170724685719 | 4.05289650832201 | 9.11511230390607E-27 | 4.23852722131632E-25 |
| aal-miR-275-5p | -4.44813211067005 | 13.7776697012477 | 1.17714531116766E-24 | 4.37898055754369E-23 |
| novel_11 | 3.82833941201173 | 3.52463339468505 | 2.75931957496839E-22 | 8.55389068240202E-21 |
| aal-miR-125-3p | -4.26494659531624 | 7.62210686236575 | 5.73542436182421E-22 | 1.52398418757043E-20 |
| aal-miR-998-5p | -3.57989956334852 | 5.94417623496733 | 1.93434465744638E-21 | 4.49735132856284E-20 |
| aal-miR-308-5p | -3.73305230841969 | 11.7640529530044 | 6.63993184196951E-20 | 1.3722525806737E-18 |
| aal-miR-79-5p | -3.77553384133528 | 8.71542058507101 | 1.50219161050041E-18 | 2.79407639553077E-17 |
| aal-miR-277-5p | -3.87752189960607 | 8.23581842333128 | 5.98756788620986E-17 | 1.01244329712276E-15 |
| aal-novel-5-5p | 2.92534802868388 | 6.15176034205506 | 3.15823738341222E-16 | 4.89526794428894E-15 |
| aal-miR-2941-5p | -4.70972580773228 | 1.93370861027017 | 3.20929766716886E-15 | 4.59176435456467E-14 |
| novel_62 | 3.04079425467027 | 6.89234878730269 | 1.06487455493276E-14 | 1.41476190869639E-13 |
| aal-miR-252-3p | -3.1183268380957 | 8.81262004914743 | 4.02522151870908E-14 | 4.99127468319926E-13 |
| novel_56 | 2.88455562644694 | 8.87439982221377 | 4.55587196711692E-13 | 5.29620116177342E-12 |
| novel_12 | 2.82940647917435 | 8.72782992524712 | 2.05116306593302E-12 | 2.2442137074326E-11 |
| aal-let-7-3p | -2.87444933954981 | 6.47771218178983 | 8.79801369019113E-12 | 9.0912808131975E-11 |
| novel_80 | -3.67897023745288 | 1.68140232212531 | 1.05374322053414E-11 | 1.03155915273342E-10 |
| aal-novel-10-5p | -5.18185910623664 | 1.15138172139682 | 1.62189763595315E-11 | 1.50836480143643E-10 |
| aal-miR-1-3p | 3.1105775211911 | 6.63637278801408 | 1.75463738664674E-11 | 1.55410739960139E-10 |
| aal-miR-7-5p | -2.29347968072914 | 12.5645174189533 | 2.63554404794081E-09 | 2.22823269507723E-08 |
| aal-miR-263a-5p | -2.59768537049161 | 5.09955785291336 | 6.38846841177771E-09 | 5.16632662865502E-08 |
| novel_86 | -3.01180797504256 | 1.71321310962084 | 8.31226071613564E-09 | 6.44200205500512E-08 |
| aal-miR-2945-5p | -2.20964568473124 | 4.85998557514981 | 1.12365588517852E-08 | 8.35999978572818E-08 |
| novel_55 | 1.84953710191012 | 6.19444580384127 | 3.27162143623597E-08 | 2.34046764284573E-07 |
| novel_71 | -2.41438757760136 | 2.20150426370704 | 4.98731310237201E-08 | 3.43570458163405E-07 |
| novel_90 | -3.00046481128921 | 1.36347206132402 | 1.39658146725436E-07 | 9.27729117533256E-07 |
| novel_101 | -2.51003032850359 | 1.93069656731235 | 1.48522712939396E-07 | 9.52593951956127E-07 |
| aal-novel-9-3p | -2.33413722282348 | 2.27064205184178 | 2.54860528441074E-07 | 1.58013527633466E-06 |
| novel_87 | -2.68528217109949 | 1.10984829938519 | 3.44630578908349E-07 | 2.0677834734501E-06 |
| aal-miR-new13 | 3.1745624037903 | 3.47231622510323 | 9.67573164109226E-07 | 5.62401901638488E-06 |
| aal-miR-9c-5p | -1.92926050887651 | 10.7224471292005 | 1.32853855664034E-06 | 7.48812641015462E-06 |
| aal-miR-279-5p | -1.57533179569277 | 4.92842879765245 | 1.50847159721829E-06 | 8.07053430590607E-06 |
| aal-miR-275-3p | 1.99596383214863 | 13.6941971844771 | 1.51864892853071E-06 | 8.07053430590607E-06 |
| novel_60 | 1.64545401119037 | 3.70083048991896 | 1.77248564331257E-06 | 9.1578424904483E-06 |
| novel_91 | -1.58416018485724 | 3.2917059485268 | 3.25589108942571E-06 | 1.63674525035995E-05 |

| | | | | |
|---|---|---|---|---|
| novel_17 | -6.41886221962339 | 0.262838845110904 | 4.20226035541615E-06 | 2.05689585817738E-05 |
| aal-novel-2-5p | -2.0852177783496 | 7.15888066633579 | 4.81234257975825E-06 | 2.29511723034624E-05 |
| novel_48 | 1.78357375381477 | 7.74713482635069 | 5.35214964507745E-06 | 2.48874958496102E-05 |
| aal-miR-125-5p | 1.60072546999516 | 10.6590330538673 | 6.77431585066951E-06 | 3.07322621518178E-05 |
| aal-miR-9b-5p | -1.80835477464594 | 9.76063050450174 | 8.53496624399151E-06 | 3.77977076519624E-05 |
| novel_49 | 1.47335619546824 | 3.52852852231895 | 9.85935657709735E-06 | 4.26474493800025E-05 |
| aal-miR-282-5p | 1.87959748191578 | 1.03702959834519 | 1.37836769950785E-05 | 5.82673618428317E-05 |
| novel_72 | 1.61795956513994 | 2.61995478349624 | 1.53279284875611E-05 | 6.33554377485861E-05 |
| aal-miR-278-3p | -1.85151064240302 | 8.74648457206226 | 3.35629567566693E-05 | 0.000135711086016 |
| aal-miR-10-5p | 1.49239459394872 | 5.64034729589154 | 3.66047075071414E-05 | 0.000144861182901 |
| aal-miR-1000-3p | -2.12804353574755 | 1.42366200060884 | 4.85216661238346E-05 | 0.00018802145623 |
| aal-miR-285-3p | 1.60392679183686 | 2.89431906444502 | 5.24953007497196E-05 | 0.000199267876315 |
| aal-miR-2765-5p | -1.577676557123 | 9.99150727860771 | 5.88148452243703E-05 | 0.000218791224235 |
| novel_18 | -1.53467389815 | 2.28584319894261 | 6.19220414355419E-05 | 0.000225833327588 |
| novel_63 | 1.36864434354537 | 5.72255722023208 | 7.04923216790697E-05 | 0.00025214561216 |
| novel_41 | 1.51084800844243 | 8.5953760844073 | 0.00010442346956 | 0.000366467270533 |
| aal-bantam-5p | -1.56549864460101 | 10.7730807667517 | 0.00011370902237 | 0.000391664410386 |
| novel_19 | -1.22786233317246 | 4.01942405947962 | 0.000137910225417 | 0.000460360498465 |
| novel_106 | 2.09987061481339 | 0.430757440884428 | 0.000138603160828 | 0.000460360498465 |
| novel_82 | -5.99650321240092 | -0.032245687816617 | 0.000158429187317 | 0.000516979453349 |
| novel_47 | 1.29686359521972 | 5.3180549821286 | 0.000161656077418 | 0.000518414317236 |
| novel_42 | 1.22671869811896 | 5.44822046757537 | 0.000171931849093 | 0.000542022439513 |
| novel_50 | 1.25503489301347 | 5.27705139431807 | 0.000186035685396 | 0.000576710624728 |
| novel_20 | 1.50638444116217 | 2.22783884767167 | 0.000356511212777 | 0.001087066976665 |
| novel_8 | 1.84701541870086 | 0.940755290837084 | 0.000372009286932 | 0.001116027860795 |
| aal-miR-87-5p | -1.13793605956629 | 3.42161170472025 | 0.000475854074842 | 0.001404902506676 |
| novel_109 | 1.46082658974475 | 5.84650266016604 | 0.000595504784905 | 0.001730685781129 |
| aal-miR-190-5p | -1.35679733933389 | 10.6178403646316 | 0.000689633100367 | 0.001973411641049 |
| aal-miR-970-3p | 1.26537242106438 | 9.99467749935239 | 0.000772804941704 | 0.002177904835711 |
| novel_84 | -2.59708840296999 | 0.29287211806864 | 0.000790393351658 | 0.002194226319529 |
| novel_46 | 1.24092702093432 | 2.75159071453278 | 0.000934245279269 | 0.00252340659273 |
| novel_25 | 2.41099449248027 | 9.8182713100982 | 0.00093610244569 | 0.00252340659273 |
| aal-novel-8-3p | -1.47596569741976 | 1.34560124880347 | 0.000955668684728 | 0.002539348219419 |
| novel_24 | 2.39244562947808 | 9.66670590011648 | 0.001178789605405 | 0.003088096712751 |
| novel_110 | 1.0572375854664 | 2.39088510473128 | 0.001551845452396 | 0.003974130410198 |
| novel_53 | 1.32297760001023 | 8.22084335823845 | 0.00155973935454 | 0.003974130410198 |
| aal-miR-new1 | -1.74690454088274 | 3.28321116762776 | 0.001750909169163 | 0.004400933857627 |
| aal-miR-970-5p | -1.75963981051303 | 1.17079067045452 | 0.00234931873969 | 0.005826310474431 |
| novel_44 | -1.25271707371148 | 13.6934018502242 | 0.002499237774901 | 0.006116555606994 |

| | | | | |
|---|---|---|---|---|
| novel_3 | 2.85875334290583 | -0.554213956421804 | 0.002562983078688 | 0.006191101982286 |
| novel_98 | -5.28585242371874 | -0.49370332316279 | 0.003022680771933 | 0.007207931071533 |
| aal-miR-new13* | 1.40611562996644 | 6.30164908865784 | 0.00338217932401 | 0.007963105750201 |
| aal-miR-92b-5p | -1.17020623075685 | 6.60892713103426 | 0.003434835261408 | 0.007985991982774 |
| aal-miR-1-5p | -2.73569852968029 | 0.001982234298544 | 0.003709300908591 | 0.008517653938246 |
| aal-miR-14-5p | -1.04046257784152 | 4.99124278325411 | 0.003802114763973 | 0.008624309098769 |
| novel_32 | 1.24235767705288 | 5.06290185927863 | 0.00773201679087 | 0.017327170157854 |
| novel_70 | -1.61813739665843 | 0.95918008632307 | 0.00785117320709 | 0.017384740672842 |
| novel_85 | -1.34929363257812 | 0.764034486953195 | 0.008378041303524 | 0.018333125675946 |
| novel_13 | 1.59066971339634 | 0.567915754466653 | 0.009412171960438 | 0.020356557960946 |
| novel_102 | 0.796038970749807 | 4.09089062681308 | 0.01148047386175 | 0.024544461359603 |
| aal-miR-999-5p | -2.44220107780331 | -0.18416487127785 | 0.01276299799167 | 0.026976336664212 |
| aal-miR-14-3p | 0.926463516682058 | 14.7426172346806 | 0.014421775433836 | 0.030139890232512 |
| novel_65 | 1.17011325014923 | 12.8583780727724 | 0.015085416196583 | 0.031176526806271 |
| novel_61 | 1.35908498615545 | 6.44959495559836 | 0.015951611690631 | 0.032604393125905 |
| novel_64 | 0.899515815838347 | 10.7498291925264 | 0.01647800642126 | 0.033314230373417 |
| aal-miR-981-3p | -1.20942899817323 | 0.838712313727188 | 0.017209183120627 | 0.034418366241253 |
| aal-miR-100-5p | 0.924541099347681 | 15.4729912038021 | 0.017468043208026 | 0.034564425922264 |
| novel_83 | 1.64660350012979 | 0.412036399548397 | 0.018452918583676 | 0.036128872174355 |
| aal-miR-965-3p | 0.80049128839996 | 6.27168184296521 | 0.02404728343674 | 0.046591611658684 |
| aal-miR-307-5p | -1.09389026850353 | 1.8125009953288 | 0.026319220344311 | 0.050467783340637 |
| novel_76 | -1.03012289766972 | 1.86698562214489 | 0.030907163334113 | 0.058153678204749 |
| aal-miR-2946-3p | -0.695683606849552 | 3.35954550164571 | 0.030952764205754 | 0.058153678204749 |
| novel_94 | 0.673407778187646 | 3.6726478947525 | 0.032497888414844 | 0.060446072451609 |
| novel_26 | 1.03579348143513 | 2.70429341183385 | 0.034013480556168 | 0.062638686964825 |
| novel_15 | -1.49403748010072 | 0.303656062094196 | 0.046582788337254 | 0.084945084614993 |

**Supplemental Table 3: List of primers used in this study**

| Primer | Sequence |
|---|---|
| aal_novel_98 | TGAAATAATACATATAGCATCC |
| aal_novel_86 | CTTAACAGATTGAAAGCTGAAA |
| aal_novel_71 | ACTATTGTAGTAGCGATAAGATG |
| aal_novel_62 | TTGATACGCTTGATACTCCTCC |
| aal_novel_11 | TCGGATAACAGCATTCGGTTGA |
| aal_miR-998-5p | TAGCACCATGAGATTCAGCTC |
| aal_miR-989-5p | GTGTGCTTTGTGACAATGAGAT |

**Supplemental Table 4: List of GO terms of target genes of the differentially expressed miRNAs**

| source | term name | term id | adjusted p-value | -log10 adjusted p-value |
|---|---|---|---|---|
| GO:Molecular Function | protein binding | GO:0005515 | 1.97E-07 | 6.70479174603375 |
| GO:Molecular Function | sequence-specific DNA binding | GO:0043565 | 9.94889369300971E-06 | 5.00222520967977 |
| GO:Molecular Function | anion binding | GO:0043168 | 0.000142492380265 | 3.84620835879342 |
| GO:Molecular Function | purine ribonucleotide binding | GO:0032555 | 0.000430151352924 | 3.36637870680591 |
| GO:Molecular Function | purine ribonucleoside triphosphate binding | GO:0035639 | 0.000443729811125 | 3.35288139307269 |
| GO:Molecular Function | purine nucleotide binding | GO:0017076 | 0.000552793955253 | 3.2574367145752 |
| GO:Molecular Function | ribonucleotide binding | GO:0032553 | 0.000615818043821 | 3.21054759016977 |
| GO:Molecular Function | transcription regulator activity | GO:0140110 | 0.001738746781564 | 2.75976366088207 |
| GO:Molecular Function | DNA-binding transcription factor activity | GO:0003700 | 0.002501260985401 | 2.60184099095407 |
| GO:Molecular Function | guanyl ribonucleotide binding | GO:0032561 | 0.004447729803985 | 2.35186160366575 |
| GO:Molecular Function | guanyl nucleotide binding | GO:0019001 | 0.005383072419859 | 2.26896977746956 |
| GO:Molecular Function | enzyme binding | GO:0019899 | 0.006181084233802 | 2.20893533794413 |
| GO:Molecular Function | purine ribonucleoside binding | GO:0032550 | 0.007851308256477 | 2.10505797112358 |
| GO:Molecular Function | purine nucleoside binding | GO:0001883 | 0.007851308256477 | 2.10505797112358 |
| GO:Molecular Function | GTP binding | GO:0005525 | 0.007851308256477 | 2.10505797112358 |
| GO:Molecular Function | lipid binding | GO:0008289 | 0.008047719337836 | 2.09432717793362 |
| GO:Molecular Function | kinase activity | GO:0016301 | 0.009089102570204 | 2.04147899555129 |
| GO:Molecular Function | cytoskeletal protein binding | GO:0008092 | 0.009478441329184 | 2.02326307382112 |
| GO:Molecular Function | nucleotide binding | GO:0000166 | 0.012114886893985 | 1.91668063613214 |
| GO:Molecular Function | nucleoside phosphate binding | GO:1901265 | 0.012114886893985 | 1.91668063613214 |
| GO:Molecular Function | ribonucleoside binding | GO:0032549 | 0.013557237634372 | 1.8678287914691 |
| GO:Molecular Function | Ran GTPase binding | GO:0008536 | 0.013964751392081 | 1.85496679143511 |
| GO:Molecular Function | nucleoside binding | GO:0001882 | 0.016175069942314 | 1.79115383273065 |
| GO:Molecular Function | microtubule binding | GO:0008017 | 0.020763762324526 | 1.6826939509807 |
| GO:Molecular Function | small molecule binding | GO:0036094 | 0.022758105400212 | 1.64286389554946 |

| | | | | |
|---|---|---|---|---|
| GO:Molecular Function | binding | GO:0005488 | 0.027749683362733 | 1.55674196802247 |
| GO:Molecular Function | kinase binding | GO:0019900 | 0.044286231717747 | 1.35373127193499 |
| GO:Molecular Function | tubulin binding | GO:0015631 | 0.048407913577793 | 1.31508363541692 |
| GO:Molecular Function | translation initiation factor binding | GO:0031369 | 0.049399835895724 | 1.30627449378283 |
| GO:Biological Process | transport | GO:0006810 | 8.51E-08 | 7.06994247853219 |
| GO:Biological Process | establishment of localization | GO:0051234 | 2.58E-07 | 6.5878507411063 |
| GO:Biological Process | localization | GO:0051179 | 8.26E-07 | 6.08307998293039 |
| GO:Biological Process | nitrogen compound transport | GO:0071705 | 2.48419778217594E-05 | 4.6048138302796 |
| GO:Biological Process | protein transport | GO:0015031 | 5.60274909074619E-05 | 4.2515988262107 |
| GO:Biological Process | peptide transport | GO:0015833 | 8.62963420463158E-05 | 4.06400761289162 |
| GO:Biological Process | amide transport | GO:0042886 | 0.000106695450036 | 3.97185410041043 |
| GO:Biological Process | organic substance transport | GO:0071702 | 0.000160156005672 | 3.79545677113447 |
| GO:Biological Process | establishment of protein localization | GO:0045184 | 0.000243286887315 | 3.61388129805698 |
| GO:Biological Process | intracellular protein transport | GO:0006886 | 0.000321347482484 | 3.49302509811086 |
| GO:Biological Process | regulation of cellular metabolic process | GO:0031323 | 0.001116734595904 | 2.95205002941786 |
| GO:Biological Process | protein localization | GO:0008104 | 0.001137107508363 | 2.94419847279533 |
| GO:Biological Process | regulation of macromolecule biosynthetic process | GO:0010556 | 0.001347068934507 | 2.8706101792509 |
| GO:Biological Process | regulation of cellular biosynthetic process | GO:0031326 | 0.001506735691224 | 2.82196292413548 |
| GO:Biological Process | regulation of biosynthetic process | GO:0009889 | 0.001506735691224 | 2.82196292413548 |
| GO:Biological Process | regulation of RNA metabolic process | GO:0051252 | 0.001742407898429 | 2.75885016890351 |
| GO:Biological Process | regulation of nitrogen compound metabolic process | GO:0051171 | 0.001944608380649 | 2.71116784690786 |
| GO:Biological Process | regulation of metabolic process | GO:0019222 | 0.002114589841486 | 2.67477385847994 |
| GO:Biological Process | regulation of nucleobase-containing compound metabolic process | GO:0019219 | 0.002183633703617 | 2.66082021113192 |
| GO:Biological Process | regulation of gene expression | GO:0010468 | 0.002185617362111 | 2.66042586803554 |
| GO:Biological Process | macromolecule localization | GO:0033036 | 0.002219267234713 | 2.65379039872639 |
| GO:Biological Process | regulation of primary metabolic process | GO:0080090 | 0.002384849514927 | 2.62253901993566 |
| GO:Biological Process | regulation of macromolecule metabolic process | GO:0060255 | 0.002566239357879 | 2.59070283862444 |
| GO:Biological Process | establishment of localization in cell | GO:0051649 | 0.002743643477944 | 2.56167232358524 |
| GO:Biological Process | phosphorylation | GO:0016310 | 0.003453481307897 | 2.4617428902874 |

| | | | | |
|---|---|---|---|---|
| GO:Biological Process | regulation of cellular macromolecule biosynthetic process | GO:2000112 | 0.003470951542013 | 2.45955144947559 |
| GO:Biological Process | cellular macromolecule localization | GO:0070727 | 0.003481604556889 | 2.45822055789234 |
| GO:Biological Process | regulation of RNA biosynthetic process | GO:2001141 | 0.004512863243424 | 2.34554782702192 |
| GO:Biological Process | regulation of nucleic acid-templated transcription | GO:1903506 | 0.004512863243424 | 2.34554782702192 |
| GO:Biological Process | cellular protein localization | GO:0034613 | 0.006185964408549 | 2.20859258307436 |
| GO:Biological Process | biological regulation | GO:0065007 | 0.007230796704623 | 2.14081384857561 |
| GO:Biological Process | cellular localization | GO:0051641 | 0.007643615822622 | 2.11670114923452 |
| GO:Biological Process | phosphorus metabolic process | GO:0006793 | 0.007684300800866 | 2.11439564308164 |
| GO:Biological Process | intracellular transport | GO:0046907 | 0.009199332664334 | 2.03624367599237 |
| GO:Biological Process | regulation of transcription, DNA templated | GO:0006355 | 0.011367291301216 | 1.94434301051739 |
| GO:Biological Process | phosphate-containing compound metabolic process | GO:0006796 | 0.012039107973232 | 1.91940569054121 |
| GO:Biological Process | regulation of biological process | GO:0050789 | 0.012141316007587 | 1.91573423716046 |
| GO:Cellular Component | plasma membrane part | GO:0044459 | 0.010635837486157 | 1.97322830723784 |
| GO:Cellular Component | cytoskeleton | GO:0005856 | 0.0287728437121 | 1.54101721327682 |
| KEGG | Glycerophospholipid metabolism | KEGG:00564 | 0.00078674336832 | 3.10416690918926 |
| KEGG | Protein processing in endoplasmic reticulum | KEGG:04141 | 0.013746309762266 | 1.86181387383253 |
